# Curcumin Enhances Alcoholic Fermentation in Yeast by Promoting Osmotic Stress Adaptation

**DOI:** 10.64898/2026.09.10.750560

**Authors:** Yukiko Nakase, Sora Kim, Keita Sakaue, Yukiko Sugimoto, Taisuke Seike, Mutsumi Watanabe, Takayuki Tohge, Yuichi Morozumi, Daisuke Watanabe

## Abstract

Spices contain diverse plant secondary metabolites with antimicrobial and antioxidant activities that can influence plant–microbe interactions. Among these metabolites, curcumin and piperine are major bioactive constituents of turmeric and black pepper, respectively. In this study, we demonstrate that curcumin and piperine enhance alcoholic fermentation in the budding yeast *Saccharomyces cerevisiae*. Curcumin significantly promoted alcoholic fermentation under high-glucose conditions (20–40% glucose), which impose substantial osmotic stress on yeast cells. Under these conditions, curcumin enhanced yeast growth, improved osmotic stress tolerance, and increased resistance to cell wall digestion, suggesting enhanced stress adaptation during fermentation. Furthermore, deletion of *HOG1*, which encodes the central MAP kinase of the High-Osmolarity Glycerol (HOG) pathway responsible for the osmotic stress response, abolished the fermentation-enhancing effect of curcumin, indicating that Hog1 function is required for this effect. Consistent with this result, transcriptomic and RT-qPCR analyses revealed increased expression of stress adaptation-related genes, including known HOG pathway-responsive genes. In contrast, the expression of genes involved in amino acid catabolism was decreased at 12 h after the initiation of fermentation, while metabolomic analysis revealed increased intracellular levels of amino acids, including glutamate and asparagine, at 24 h. These coordinated transcriptional and metabolic changes are consistent with reduced amino acid catabolism and a shift toward a metabolic state favorable for sustained fermentation. Together, these findings suggest that curcumin enhances alcoholic fermentation under osmotic stress by promoting stress adaptation that supports yeast growth and sustained fermentation.

**Importance:** Plant secondary metabolites are widely recognized for their roles in plant defense, whereas their beneficial effects on microbial physiology and metabolism remain poorly understood. This study demonstrates that the spice-derived compounds curcumin and piperine enhance alcoholic fermentation in the budding yeast *Saccharomyces cerevisiae*. Curcumin, in particular, improved yeast growth and stress tolerance under high-glucose conditions that impose substantial osmotic stress. Its fermentation-enhancing effect required Hog1 function and was accompanied by increased expression of HOG pathway-responsive genes and changes in amino acid metabolism. These findings demonstrate that plant-derived compounds can not only exert antimicrobial effects but also beneficially modulate yeast stress adaptation and fermentation performance. The use of natural bioactive compounds to improve fermentation robustness under high-sugar conditions may contribute to increased productivity and reduced costs in fermented food and bioethanol production.

## Introduction

Alcoholic fermentation in yeast is a crucial metabolic pathway that allows cells to survive under anaerobic or hypoxic conditions. Through glycolysis, glucose is converted into ethanol and carbon dioxide. Although fermentation yields less ATP than aerobic respiration, it does not require oxygen and thus offers a competitive advantage in low-oxygen environments. Moreover, ethanol, one of the major fermentation products, has antimicrobial properties that suppress the growth of competing microorganisms, thereby contributing to the ecological dominance of yeast in fermentative environments (1). Beyond its biological significance, alcoholic fermentation is central to a wide range of industrial processes, including the production of alcoholic beverages such as beer, wine, and sake, as well as leavened bread, where it enhances flavor and texture (2). In addition, it plays an important role in the production of bioethanol, a renewable alternative to fossil fuels that is expected to contribute to resource sustainability and environmental protection (3).

Extensive biochemical and genetic studies have elucidated the enzymes and genes involved in glycolysis and fermentation in *Saccharomyces cerevisiae* (4–6). However, direct enhancement of fermentative capacity through modification of these known genes remains limited. Even when key glycolytic and fermentative enzymes are overexpressed in laboratory strains, the high fermentation performance of sake yeast strains cannot be fully reproduced (7). These findings suggest that, despite the identification of individual enzyme functions, the regulatory mechanisms that control fermentation in yeast and their impact on fermentation performance are still not fully understood.

Meanwhile, certain plant-derived compounds have been reported to influence fermentation. For example, iso-α-acids found in hops not only impart bitterness to beer but also exert antimicrobial activity that suppresses bacterial contamination during fermentation (8). Capsaicin, the pungent compound in chili peppers, has also been shown to inhibit undesirable microorganisms and improve the preservation of fermented foods (9). Understanding how plant-derived compounds affect microorganisms, particularly yeast, is of great importance for food preservation, fermentation technology, and the development of functional foods. However, the specific mechanisms by which such compounds act during fermentation remain largely unexplored.

In this study, we investigated the effects of spice-derived plant metabolites on yeast alcoholic fermentation. Spices have long been used for their preservative and antimicrobial properties in both food and medicine. The alkaloids and terpenoids in spices are thought to have evolved as chemical defenses against microbial invasion (10).

Nevertheless, the full biological significance of these compounds is not well understood, and studies on their antimicrobial mechanisms are still limited. While most previous assessments have used microbial growth as a readout of antimicrobial activity, we focused on alcoholic fermentation—a key energy metabolism in *S. cerevisiae*—as an alternative functional marker to uncover novel effects of spice compounds. Surprisingly, while many spice components suppressed fermentation, we found that two compounds— curcumin derived from turmeric and piperine from black pepper—significantly enhanced alcoholic fermentation in yeast. This unexpected discovery suggests a novel mode of action distinct from the conventional preservative or antimicrobial function of spices. Among them, curcumin exhibited the most pronounced effect, increasing fermentation activity by approximately threefold. Further analyses suggested that curcumin enhances alcoholic fermentation by improving yeast adaptation to high-glucose conditions, thereby supporting cell proliferation and sustained fermentation. This effect was accompanied by enhanced expression of genes associated with stress adaptation and coordinated changes in amino acid metabolism. Together, these findings identify spice-derived metabolites as previously unrecognized modulators of yeast fermentation performance and provide a new perspective on how plant-derived compounds can beneficially influence microbial physiology and metabolism under industrially relevant conditions.

## Results

### Curcumin and piperine promote alcoholic fermentation

To explore novel effects of spice-derived compounds, we investigated the impact of 11 representative phytochemicals on alcoholic fermentation in *Saccharomyces cerevisiae*. Fermentation activity was assessed by measuring the amount of carbon dioxide produced during the fermentation process. The results revealed that these compounds exerted markedly different effects on alcoholic fermentation (Supplementary Fig. S1). Specifically, linalool (a major component of lavender oil), cuminaldehyde (a flavor compound in cumin), limonene (a citrus-derived terpene), allyl isothiocyanate (a pungent compound in wasabi), gingerol (a pungent component of ginger), and allicin (a sulfur-containing compound from garlic) all exhibited inhibitory effects on alcoholic fermentation. This suppression is likely attributable to the known antimicrobial properties of these compounds. In contrast, menthol (from mint), capsaicin (from chili pepper), and dipotassium glycyrrhizinate (a sweet component of licorice) exhibited no significant impact on fermentation, suggesting that their antimicrobial activity does not notably affect yeast under the tested conditions. Interestingly, curcumin (a pigment from turmeric) and piperine (a pungent compound from black pepper) markedly enhanced alcoholic fermentation (Fig. 1; Supplemental Fig. S1). A comparison of fermentation rates at 24 hours (1 day) revealed a 1.7-fold increase with piperine and an approximately 3.0-fold increase with curcumin, compared to the untreated control (Fig. 1A). Consistently, ethanol concentrations at 48 hours (2 days) were elevated by approximately 1.5-fold with piperine and approximately 2.0-fold with curcumin relative to the control (Fig. 1B). These results clearly demonstrate that curcumin and piperine exert a stimulatory effect on yeast alcoholic fermentation.

**Figure 1.**
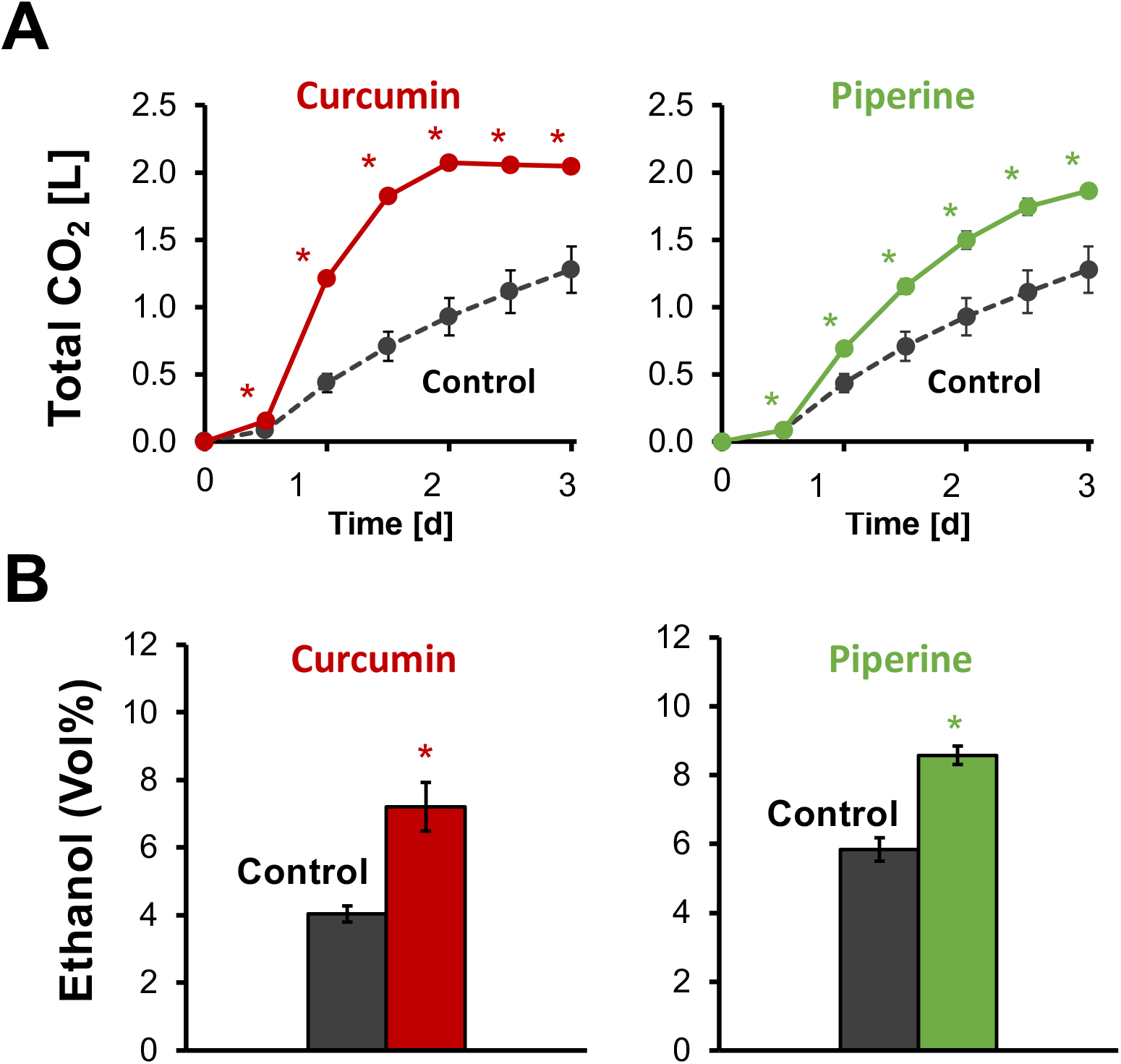
Curcumin and piperine promote alcoholic fermentation. (A) Cumulative CO₂ production by *Saccharomyces cerevisiae* X2180 measured using a Fermograph-based fermentation assay. Cells were fermented in YPD20 medium, with or without curcumin (1 mg/mL) or piperine (1 mg/mL). Cultures were incubated at 30°C for 3 days under static conditions. CO₂ production was measured at the indicated time points. (B) Ethanol concentrations in cultures of strain X2180 after 2 days of fermentation under the same conditions as in panel A. Ethanol concentrations are expressed as % (v/v). Data represent the mean ± standard deviation (SD) from three independent biological experiments. An asterisk indicates a significant difference from the corresponding untreated control (Student’s *t-test*, P < 0.05).

### Curcumin enhances alcoholic fermentation across diverse yeast strains

Curcumin, a lipophilic polyphenol predominantly found in turmeric, is well known for its anti-inflammatory and antioxidant activities. It also exhibits antimicrobial effects against bacteria, viruses, and fungi, and has attracted attention due to its broad physiological properties (11, 12). However, limited information is available regarding its effects on microbial metabolism. In our study, we found that curcumin, contrary to its well-known antimicrobial role, significantly enhanced alcoholic fermentation in yeast (Fig. 1). Among the tested compounds, curcumin exhibited the most potent stimulatory effect, exceeding that of piperine. Based on these findings, we focused subsequent analyses on elucidating the mechanism underlying curcumin’s action.

To determine whether curcumin’s fermentation-promoting effect was dose-dependent, we conducted fermentation assays using curcumin at concentrations ranging from 0.01 to 10 mg/ml. A clear enhancement of fermentation was observed from as low as 0.01 mg/ml, and the effect plateaued between 0.1 and 10 mg/ml, indicating that the stimulatory effect was not strongly dose-sensitive within this range (Supplemental Fig. S2).

Next, we evaluated whether curcumin’s effect was strain specific. In addition to the laboratory strain X2180 used in Fig. 1, we tested six additional yeast strains with diverse genetic backgrounds commonly used in industrial and academic contexts. These included the sake yeast strain K701, the widely studied laboratory strains S288C and its derivative BY4741, the laboratory strain Σ1278b, the haploid derivative X2180-1A, and the S288C-related strain BMA64-1A. Fermentation assays revealed that curcumin enhanced alcoholic fermentation across all tested strains (Fig. 2A–G). Notably, the effect was also observed in the highly fermentative sake yeast K701, indicating that curcumin is effective even in strains with strong inherent fermentation capacity.

**Figure 2.**
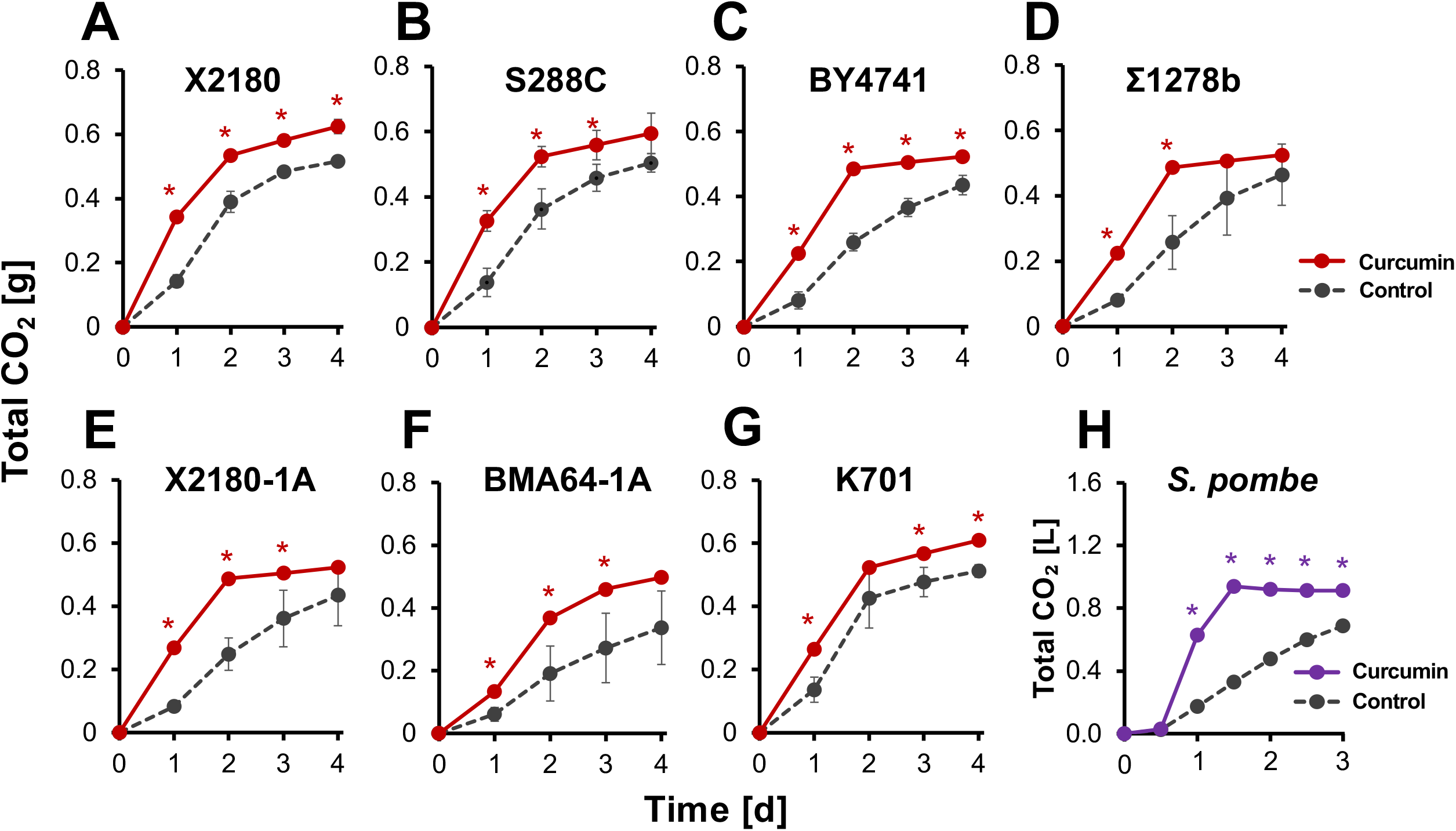
Curcumin promotes alcoholic fermentation in diverse yeast strains. Cumulative CO₂ production by the *Saccharomyces cerevisiae* laboratory strains X2180 (A), S288C (B), BY4741 (C), Σ1278b (D), X2180-1A (E), and BMA64-1A (F), the sake yeast strain K701 (G), and the *Schizosaccharomyces pombe* laboratory strain L972 (H). CO₂ production was measured using a tube-based fermentation assay for panels A–G and a Fermograph-based assay for panel H, as described in Materials and Methods. Cells were fermented statically in YPD20 medium with or without curcumin (1 mg/mL) at 30°C for 4 days. Data represent the means ± standard deviations (SD) from three independent biological experiments. An asterisk indicates a significant difference from the untreated control at the corresponding time point (Student’s *t* test, *P* < 0.05).

We further extended our analysis to the evolutionarily distant fission yeast *Schizosaccharomyces pombe*, which is also capable of alcoholic fermentation (13, 14).

Remarkably, curcumin significantly enhanced fermentation in *S. pombe* as well (Fig. 2H), suggesting that its effect transcends evolutionary divergence within fungi.

Collectively, these results demonstrate that curcumin promotes alcoholic fermentation across a wide range of yeast strains, including both budding and fission yeasts. This suggests that its fermentation-enhancing activity is not strain-specific but may instead reflect a broadly conserved physiological response.

### Curcumin analogs and mixtures promote alcoholic fermentation

Curcumin exhibits intrinsic fluorescence owing to its conjugated structure, with an absorption maximum at approximately 430 nm. Taking advantage of its fluorescence, which can be detected using a fluorescein isothiocyanate (FITC) filter set, we examined the distribution of curcumin in yeast cells during fermentation. Curcumin-derived fluorescence was concentrated around the cell periphery, giving the appearance that curcumin surrounded the cells (Fig. 3A). We next examined whether structurally related compounds also enhance alcoholic fermentation. Turmeric naturally contains the curcuminoids desmethoxycurcumin and bisdemethoxycurcumin. We also examined tetrahydrocurcumin, a reduced metabolite of curcumin formed during intestinal absorption (15, 16). Although these analogs differ in the number of conjugated double bonds and specific functional groups, their overall structures resemble that of curcumin (Fig. 3B), and each has been reported to exhibit antimicrobial and antioxidant properties (17–19). To determine whether these curcumin analogs also affect yeast alcoholic fermentation, we evaluated their effects in fermentation assays. All tested curcuminoids were found to promote alcoholic fermentation, similar to curcumin itself (Fig. 3B). Curcumin is known to undergo non-enzymatic degradation, producing breakdown products such as vanillin and ferulic acid via cleavage of the central carbon chain linking the phenolic rings (20). We therefore examined the effects of these degradation products on yeast fermentation. In contrast to curcumin and its analogs, both vanillin and ferulic acid suppressed alcoholic fermentation (Fig. 3C). These phenolic compounds have been reported to destabilize cellular membranes and inhibit protein synthesis, leading to cytotoxicity and impaired yeast growth (21, 22). Our findings are consistent with these reports, suggesting that reduced metabolic activity due to cellular stress likely caused the observed inhibition of fermentation. Taken together, these results suggest that the fermentation-enhancing activity is associated with curcumin and its structurally related analogs rather than with its degradation products.

**Figure 3.**
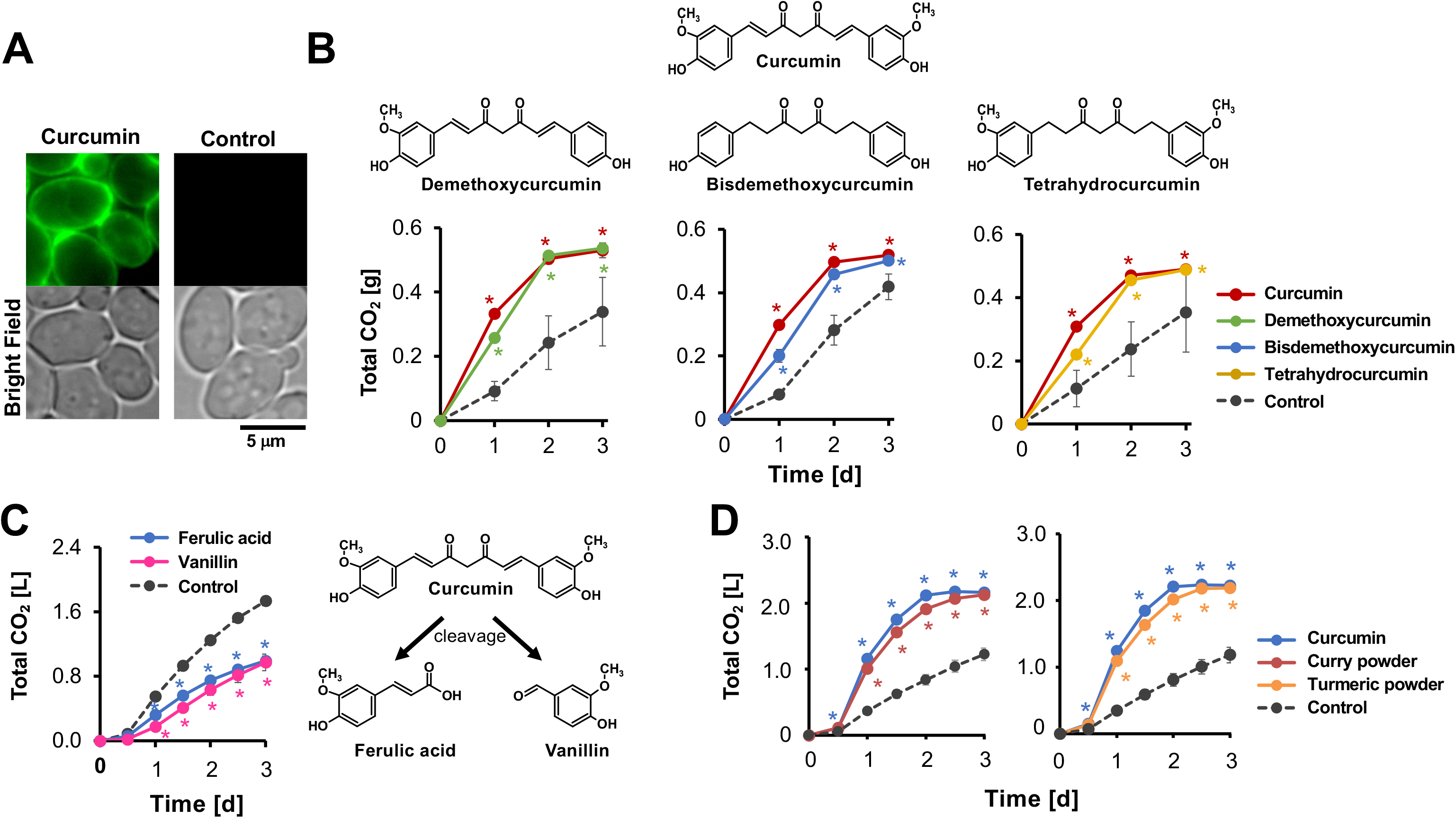
Effect of curcumin analogs and mixtures on alcoholic fermentation. (A) Representative bright-field and fluorescence images of X2180 cells fermented in YPD20 medium, with or without curcumin (1 mg/mL), at 30°C under static conditions. Curcumin-associated autofluorescence was observed by fluorescence microscopy after 24 hr of fermentation. Scale bar, 5 µm. (B) Effects of curcumin and its analogs on alcoholic fermentation. Cumulative CO₂ production by strain X2180 in the absence or presence of curcumin, demethoxycurcumin, bisdemethoxycurcumin, or tetrahydrocurcumin (1 mg/mL each) was measured using a tube-based fermentation assay. CO₂ production was estimated from the decrease in culture weight. (C) Effects of the curcumin degradation products ferulic acid and vanillin on alcoholic fermentation. Cumulative CO₂ production by strain X2180 in the absence or presence of ferulic acid or vanillin (1 mg/mL each) was measured using a Fermograph-based fermentation assay. (D) Effects of commercial curry powder and turmeric powder on alcoholic fermentation. Cumulative CO₂ production by strain X2180 in the absence or presence of curcumin, commercial curry powder, or commercial turmeric powder (1 mg/mL each) was measured using a Fermograph-based fermentation assay. Unless otherwise indicated, fermentation was performed in YPD20 medium at 30°C for 3 days under static conditions, and CO₂ production was measured at the indicated time points. Data in panels B–D represent the means ± standard deviations (SD) from three independent biological experiments. An asterisk indicates a significant difference from the corresponding untreated control (Student’s *t* test, *P* < 0.05).

Thus far, we have used purified curcumin in our experiments. However, curcumin is also readily available in the form of turmeric powder, derived from the rhizome of *Curcuma longa*, and is a common ingredient in curry powder. To evaluate whether turmeric and curry powder—both of which contain curcumin—could similarly influence fermentation, we conducted assays using these crude mixtures. Notably, both turmeric powder and curry powder significantly enhanced yeast alcoholic fermentation (Fig. 3D). These results demonstrate that the fermentation-promoting effect is not limited to purified curcumin, but is also evident in curcumin-containing mixtures such as turmeric and curry powder.

### Curcumin promotes alcoholic fermentation under high-glucose conditions

In industrial fermentation processes, such as wine production and sweet bread manufacturing, the sugar content of grape juice or dough often exceeds 20%. These high-glucose environments impose severe osmotic stress on yeast cells, resulting in impaired growth and reduced fermentation capacity. To assess whether curcumin could mitigate these limitations, we evaluated its effect on alcoholic fermentation under varying high-glucose conditions. Fermentation assays were performed using media containing 10– 40% glucose. At all tested glucose concentrations, curcumin supplementation significantly enhanced alcoholic fermentation compared to the untreated control (Fig. 4). While increasing glucose concentrations led to reduced fermentation rates in control samples, curcumin-treated samples exhibited a marked improvement in fermentation activity. Under 10% glucose—a condition with minimal osmotic stress—fermentation proceeded rapidly in the early phase. However, in the curcumin-treated group, glucose was depleted within 24 hours, resulting in an early termination of fermentation due to substrate exhaustion. These results indicate that curcumin helps maintain fermentation efficiency even under high-glucose stress, suggesting its potential to counteract osmotic inhibition and promote alcoholic fermentation under industrially relevant conditions.

**Figure 4.**
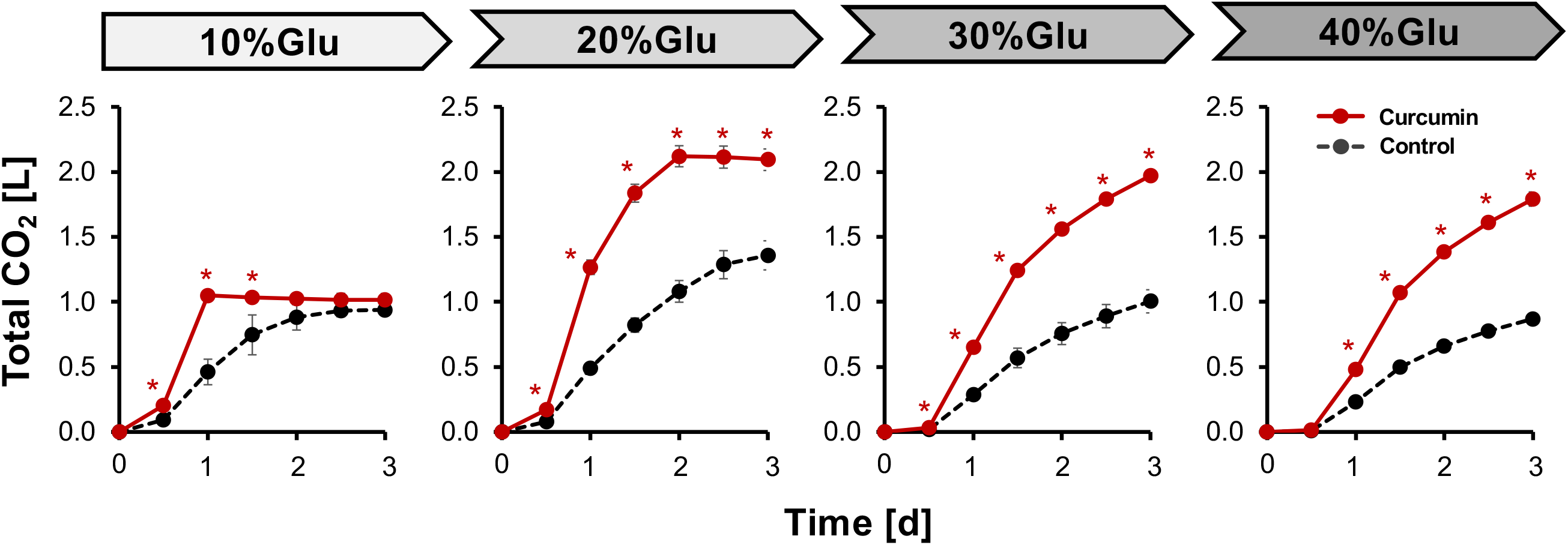
Curcumin promotes alcoholic fermentation under high-glucose conditions. Cumulative CO₂ production by strain X2180 during fermentation in YPD media containing 10%, 20%, 30%, or 40% glucose, with or without curcumin (0.1 mg/mL), was measured using a Fermograph-based fermentation assay. Cultures were incubated at 30°C for 3 days under static conditions, and CO₂ production was measured at the indicated time points. Data represent the means ± standard deviations (SD) from three independent biological experiments. An asterisk indicates a significant difference from the corresponding untreated control at the indicated time point (Student’s *t* test, *P* < 0.05).

### Curcumin improves osmotic stress tolerance

To determine whether the stimulatory effect of curcumin on alcoholic fermentation is associated with enhanced yeast growth, we measured cell proliferation under fermentative conditions. First, yeast cultures were grown in media containing either 20% or 40% glucose, with or without curcumin supplementation. As expected, high glucose concentrations significantly impaired yeast growth; however, the addition of curcumin markedly improved cell proliferation under these conditions (Fig. 5A). By day 2 of fermentation, cell densities in the curcumin-treated cultures were approximately four times higher than those in the untreated controls. These findings suggest that curcumin may accelerate ethanol production in part by increasing the viable cell population within the fermentation vessel.

**Figure 5.**
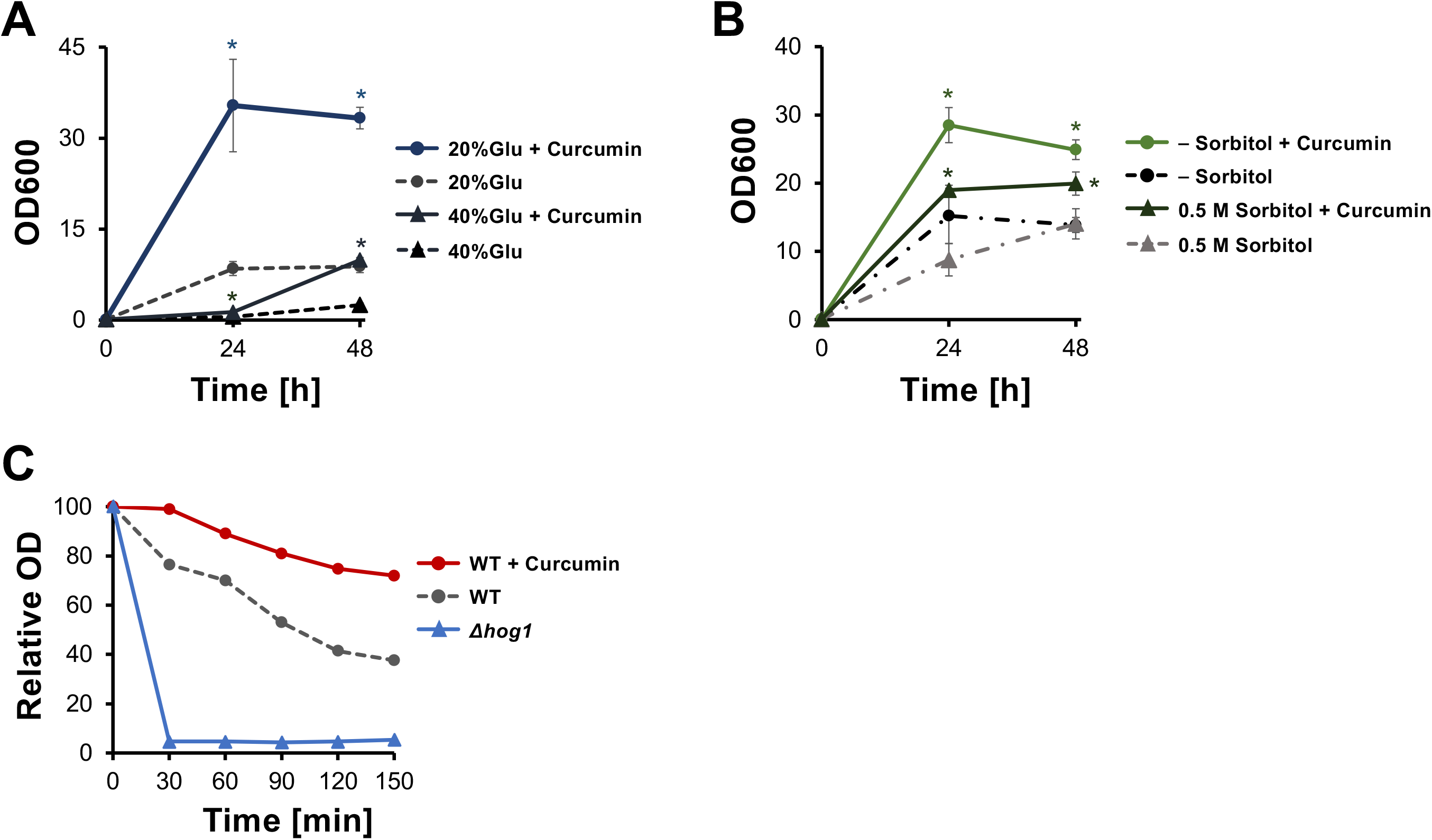
Curcumin improves osmotic stress tolerance during alcoholic fermentation. (A) Growth of strain X2180 during fermentation in YPD media containing 20% or 40% glucose, with or without curcumin (0.1 mg/mL). Cultures were incubated at 30°C under static conditions, and cell growth was monitored by measuring the optical density at 600 nm (OD₆₀₀) at the indicated time points. (B) Growth of strain X2180 in YPD medium with or without 0.5 M sorbitol and with or without curcumin (0.1 mg/mL). Cultures were incubated at 30°C under static conditions, and OD₆₀₀ was measured at the indicated time points. (C) Sensitivity of strain X2180 and the *Δhog1* mutant to Zymolyase. X2180 cells were fermented for 1 day with or without curcumin (0.1 mg/mL), collected, washed, and subsequently treated with Zymolyase. Untreated *Δhog1* cells were included as a Zymolyase-sensitive control. Cell lysis was monitored by measuring OD₆₀₀ at the indicated time points. A decrease in OD₆₀₀ indicates cell lysis. Data represent the means ± standard deviations (SD) from three independent biological experiments. An asterisk indicates a significant difference from the corresponding untreated control at the indicated time point (Student’s *t* test, *P* < 0.05).

To further evaluate curcumin’s effect on osmotic stress tolerance, we conducted stress assays using D-sorbitol, a sugar alcohol commonly employed as a non-metabolizable osmoticum. When 0.5 M D-sorbitol was added to medium containing 10% glucose, cell growth was notably inhibited in the absence of curcumin, whereas curcumin supplementation significantly alleviated this growth inhibition (Fig. 5B). These results indicate that curcumin may either mitigate osmotic stress directly or enhance the intrinsic stress resistance of yeast cells.

To examine whether curcumin treatment induces cellular changes associated with enhanced stress tolerance, we next assessed yeast tolerance to the cell wall– degrading enzyme Zymolyase (Fig. 5C). If curcumin treatment enhances cellular stress resistance, treated cells would be expected to exhibit increased resistance to Zymolyase-induced lysis. Cells were cultured for one day under fermentation conditions with or without curcumin and then washed prior to Zymolyase treatment to minimize the carryover of curcumin into the assay. Thus, the assay was designed to primarily assess changes in the cells induced during fermentation rather than a direct effect of curcumin on Zymolyase activity. After the addition of Zymolyase, OD₆₀₀ was monitored over time.

As a control for increased sensitivity to cell wall stress, we also examined *Δhog1* cells, as Hog1 plays an important role in cellular stress responses, including responses associated with cell wall integrity (23). As expected, *Δhog1* cells exhibited markedly increased sensitivity to Zymolyase, with OD₆₀₀ rapidly decreasing to substantially lower levels than those observed in untreated WT cells. In contrast, curcumin-treated WT cells showed a slower decline in OD₆₀₀ than untreated WT cells, indicating greater resistance to Zymolyase-induced lysis. These results suggest that curcumin treatment during fermentation induces cellular changes that increase resistance to cell wall–degrading stress. Together with our other findings, this increased stress tolerance may contribute to improved fermentation efficiency under high-stress conditions.

### Curcumin enhances the expression of genes associated with stress adaptation

To elucidate the molecular mechanisms underlying curcumin’s effects on yeast cells, we performed transcriptomic analysis to compare global gene expression profiles between curcumin-treated and untreated cultures 12 h after the initiation of fermentation. A total of 58 genes were significantly upregulated in the curcumin-treated group compared to the untreated group (log₂FC ≥ 2) (Fig. 6A and B; Supplementary Table S1). Gene ontology (GO) analysis revealed that the upregulated genes were highly enriched in categories related to cell wall biosynthesis and structural integrity (Fig. 6C). Notably, many of these genes are known to be induced under osmotic stress conditions (24). Yeast cells adapt to osmotic stress primarily through activation of the High-Osmolarity Glycerol (HOG) pathway, which orchestrates a variety of protective responses, including transient cell cycle arrest, induction of stress-responsive genes, and increased glycerol production (25–27). In our analysis, many of the genes upregulated by curcumin were associated with cellular stress responses, including responses regulated by the HOG pathway (Fig. 6D). Among these were several members of the seripauperin (*PAU*) gene family, a large multigene family whose expression is induced under anaerobic and other environmental stress conditions (28, 29). *DAN1*, which encodes an anaerobically induced cell wall mannoprotein related to the Pau proteins, was also upregulated (30). Curcumin treatment additionally increased the expression of *GPP2*, which encodes a glycerol-3-phosphatase involved in glycerol biosynthesis (31), and *HXT1,* which encodes a low-affinity glucose transporter (32). Furthermore, *GRE2*, encoding an oxidoreductase, and *CTT1*, encoding cytosolic catalase, were significantly upregulated (33). Collectively, these transcriptomic data suggest that curcumin enhances a broad stress-adaptive transcriptional response that includes genes involved in cell wall remodeling, glycerol metabolism, and cellular stress protection.

**Figure 6.**
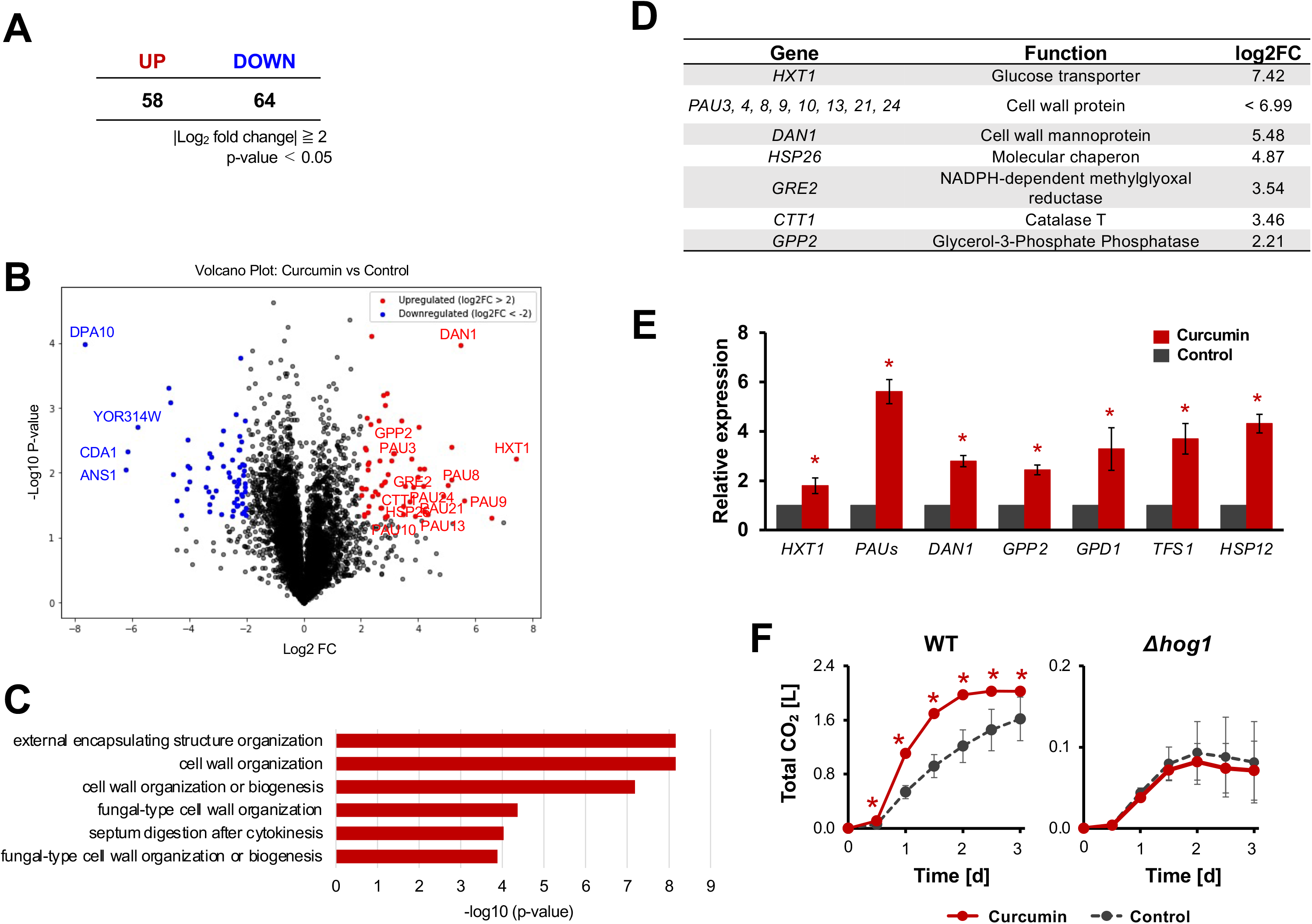
Curcumin enhances the expression of genes associated with stress adaptation during alcoholic fermentation. Strain X2180 was fermented in YPD20 medium with or without curcumin (0.1 mg/mL) at 30°C under static conditions. Cells were harvested 12 h after the initiation of fermentation and subjected to RNA-seq analysis. (A) Numbers of genes significantly upregulated or downregulated in curcumin-treated cells relative to untreated control cells (*P* < 0.05). (B) Volcano plot showing transcriptional differences between curcumin-treated and untreated cells. The *x*-axis indicates the log₂ fold change, and the *y*-axis indicates −log₁₀(*P* value). (C) Gene Ontology (GO) enrichment analysis of genes significantly upregulated in curcumin-treated cells. (D) Selected stress-responsive genes upregulated in curcumin-treated cells relative to untreated control cells. (E) RT-qPCR validation of representative stress-responsive genes identified as upregulated by RNA-seq. For RT-qPCR, RNA was extracted from independent cultures prepared separately from those used for RNA-seq and harvested 12 h after the initiation of fermentation under the same conditions. Expression levels were normalized to those of *ACT1* and are presented relative to those in untreated control cells. Data in panel E represent the means ± standard deviations (SD) from three independent biological experiments. An asterisk indicates a significant difference from the untreated control (Student’s *t* test, *P* < 0.05). (F) Cumulative CO_2_ production by strain BY4741 and the ***Δ****hog1* strain in the presence or absence of curcumin (0.1 mg/mL). Data represent the mean ± standard deviation (SD) from three independent experiments. An asterisk indicates a significant difference from the untreated control for the corresponding strain (Student’s *t* test, P < 0.05).

To independently validate the RNA-seq results and further examine the association between curcumin treatment and HOG pathway-related gene expression, we performed RT-qPCR using biological samples prepared separately from those used for RNA-seq. We analyzed four representative genes shown in Fig. 6D (*HXT1, PAU9, DAN1,* and *GPP2*), together with three established Hog1-responsive genes (*GPD1, TFS1,* and *HSP12*). All seven genes showed higher expression in curcumin-treated cells than in untreated control cells (Fig. 6E). These results independently confirmed the RNA-seq findings and were consistent with the enhancement of a HOG pathway-associated transcriptional response by curcumin. Such an enhanced stress-adaptive response may contribute to the improved osmotic stress tolerance and fermentation performance observed in curcumin-treated cells. To determine whether Hog1 is required for the fermentation-enhancing effect of curcumin, we next examined alcoholic fermentation in a *hog1* deletion strain. Curcumin enhanced fermentation in the wild-type strain, whereas this effect was abolished in the *hog1* deletion strain (Fig. 6F). These results indicate that Hog1 is required for the curcumin-mediated enhancement of alcoholic fermentation. Together, the transcriptional and genetic evidence suggests that Hog1-dependent stress adaptation contributes to the improved osmotic stress tolerance and fermentation performance of curcumin-treated cells.

### Curcumin alters amino acid metabolism during fermentation under osmotic stress conditions

Having identified the upregulation of stress adaptation-related genes in curcumin-treated cells, we next examined the genes that were downregulated in the same RNA-seq dataset. GO enrichment analysis revealed that the downregulated genes were enriched in categories associated with amino acid catabolism, including L-amino acid catabolism, aromatic amino acid catabolism, and the kynurenine metabolic pathway (Fig. 7A and Supplementary Table S2). Because these processes contribute to metabolic adaptation in response to amino acid availability and nutritional status, their downregulation suggests that curcumin alters the transcriptional regulation of amino acid metabolism during the mid-phase of fermentation. To investigate how the observed transcriptional changes are reflected in metabolic states during fermentation progression, a time-course metabolomic analysis was conducted throughout the fermentation process. Comparison of intracellular levels of the TCA cycle intermediates, citrate and 2- oxoglutarate (αKG), together with the key nitrogen metabolism–related amino acids glutamate and asparagine, showed a tendency toward higher relative levels under the untreated condition at 6 hours after the initiation of fermentation. In contrast, from the time point of 12 hours, the levels of these metabolites were elevated under the curcumin-treated condition (Fig. 7B). In particular, glutamate and asparagine were significantly increased under the curcumin-treated condition during the late phase of fermentation (24 h), showing changes that are consistent with the decrease of gene expression related to amino acid catabolism observed during the mid-phase of fermentation. Furthermore, based on the reduction in the expression of gene sets related to aromatic amino acid catabolism and the kynurenine metabolic pathway observed in Fig. 7A, intracellular levels of tryptophan and kynurenic acid were evaluated (Fig. 7C). Both metabolites exhibited time-dependent changes, with a slight increase under the curcumin-treated condition during the late phase of fermentation (24 h). These results suggest that changes in the transcriptional regulation related to aromatic amino acid catabolism may be reflected at the metabolic level. Next, comprehensive analysis of the metabolomic profiling data was conducted to explore associations with other metabolic processes. Under the curcumin-treated condition, changes in the metabolic profile, particularly in amino acid–related metabolites, were observed during the late phase of fermentation (Supplementary Fig. S3).

**Figure 7.**
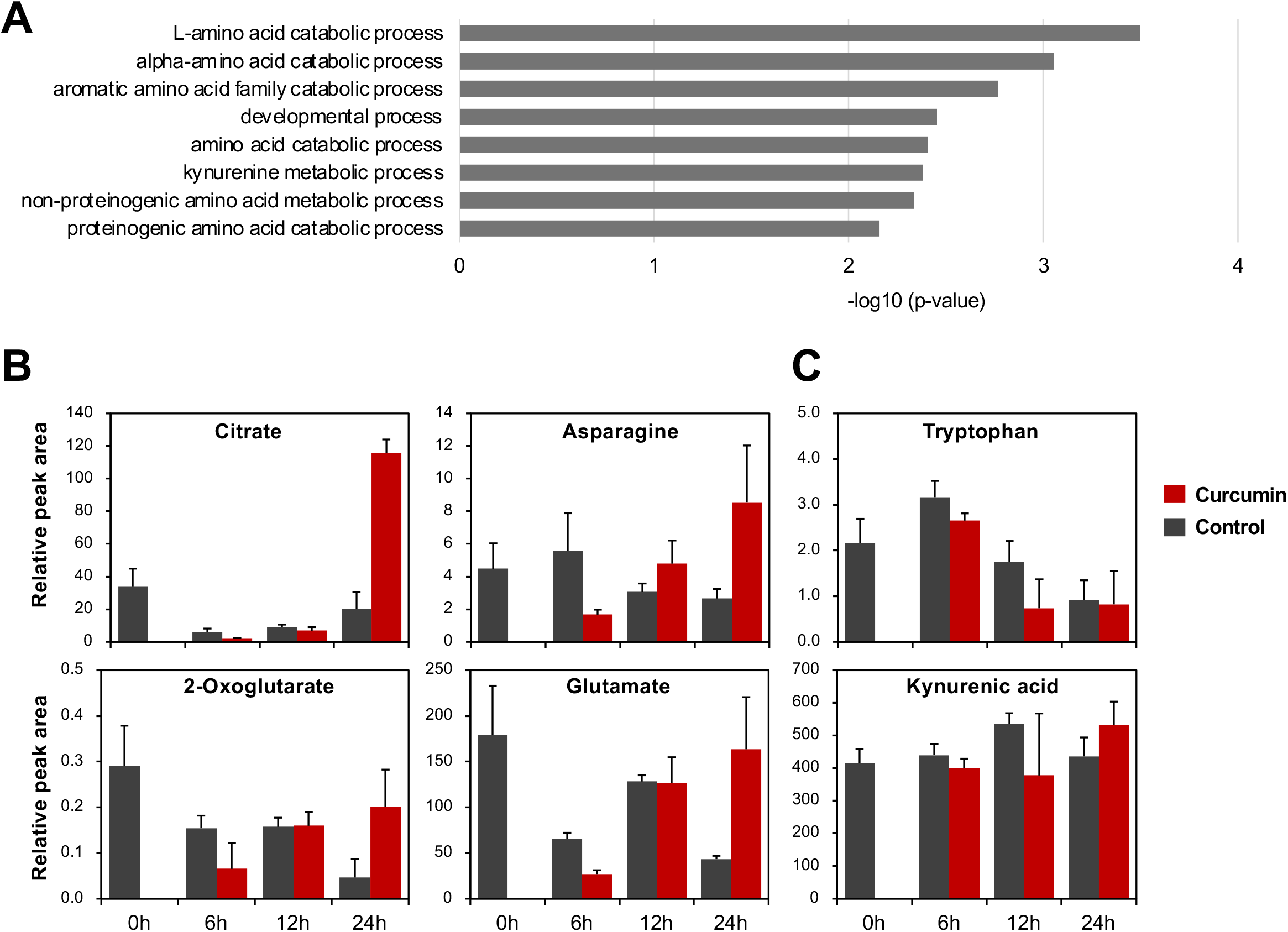
Coordinated changes in metabolic gene expression and intracellular metabolite profiles in curcumin-treated yeast. Strain X2180 was fermented in YPD20 medium with or without curcumin (0.1 mg/mL) at 30°C under static conditions. (A) GO enrichment analysis of genes significantly downregulated in curcumin-treated cells relative to untreated control cells, based on the RNA-seq data obtained 12 h after the initiation of fermentation. (B-C) Metabolite profiling by GC-MS was performed at 0, 6, 12, and 24 h in the presence or absence of curcumin. (B) TCA cycle intermediates and core amino acids, including citrate, 2- oxoglutarate, glutamate, and asparagine, were quantified. (C) Aromatic amino acids and kynurenine pathway metabolites, including tryptophan and kynurenic acid, were quantified. Metabolite levels are shown as relative abundances normalized to cell biomass. Data represent mean ± SD. Statistical significance was determined by Student’s *t* test, and detailed results are shown in Supplemental Figure S3.

Collectively, the downregulation of amino acid catabolism-related genes and the accompanying accumulation of several related metabolites suggest that curcumin alters amino acid utilization during fermentation under osmotic stress conditions. These coordinated transcriptional and metabolic changes may contribute to the enhanced fermentation observed in curcumin-treated cells.

## Discussion

Spice-derived compounds comprise diverse secondary metabolites that plants have evolved to produce. These compounds serve protective roles through their antimicrobial, antioxidant, and insect-deterrent activities, thereby helping plants defend themselves against biotic threats and environmental stress (34–36). Although these compounds have long been used for food preservation and medicinal purposes, they are also thought to function as ecological mediators of interactions between plants and other organisms, including microorganisms and animals (35). Curcumin and piperine, in particular, are known to possess antimicrobial activities, including antifungal activity against yeasts such as *Candida albicans* (11, 12, 37). Accordingly, these compounds would generally be expected to inhibit, rather than promote, yeast growth and metabolism. In contrast to this expectation, the present study demonstrated that curcumin and piperine promoted alcoholic fermentation in yeast. The increases in CO₂ release and ethanol production induced by curcumin were accompanied by enhanced yeast cell proliferation.

These findings indicate that spice-derived compounds do not invariably inhibit microbial growth and that their effects may vary depending on factors such as compound identity, concentration, and nutritional conditions. At least under the high-glucose fermentation conditions used in this study, curcumin appears to promote yeast stress adaptation and establish a physiological state favorable for maintaining cell proliferation and fermentation. Such effects are also of interest in the context of plant–microbe interactions in natural environments. Ethanol and other volatile fermentation products produced by yeast may influence ecological interactions associated with plants, including the behavior of animals that consume fruits (38, 39). Conversely, plant-derived compounds may alter the physiology and metabolism of yeasts inhabiting plant surfaces or internal tissues. However, whether these effects have ecological significance in natural environments remains to be determined.

The fermentation-enhancing effect of curcumin observed in this study appears to be associated, at least in part, with adaptation to osmotic stress under high-glucose conditions. The HOG pathway plays a central role in yeast adaptation to high-osmolarity environments. Water loss from cells following osmotic stress elicits a Hog1-mediated response that restores intracellular osmotic balance through glycerol synthesis, cell wall remodeling, and the induction of stress-responsive genes (40–42). Our RNA-seq and RT-qPCR analyses showed that curcumin treatment increased the expression of genes associated with cell wall organization and cellular stress adaptation, including several genes known to be responsive to HOG pathway signaling (Fig. 6C–E). This transcriptional response is consistent with the idea that curcumin enhances yeast adaptation to high-glucose conditions. Furthermore, the stimulatory effect of curcumin on alcoholic fermentation observed in the wild-type strain was absent in the *hog1* deletion strain (Fig. 6F), indicating that Hog1 function is involved in the fermentation-enhancing effect of curcumin. Nevertheless, the precise mechanism by which curcumin promotes fermentation through Hog1 remains unclear. Given the hydrophobicity and poor aqueous solubility of curcumin, interaction with the yeast cell surface may represent one possible mechanism. Previous encapsulation studies have shown that curcumin can associate with the plasma membrane bilayer and cell wall components of *S. cerevisiae* (43). Consistent with this report, fluorescence microscopy in the present study revealed strong curcumin-associated fluorescence at the cell periphery (Fig. 3A), supporting the possibility that curcumin interacts with the yeast cell surface. Such interactions could potentially affect membrane organization or the activity of membrane-localized stress sensors. However, whether curcumin directly affects these sensors or Hog1 signaling remains to be determined.

In addition to this stress-adaptive response, curcumin treatment was associated with changes in amino acid metabolism. GO enrichment analysis of the downregulated genes revealed significant enrichment of categories related to L-amino acid catabolism, aromatic amino acid catabolism, and the kynurenine metabolic pathway (Fig. 7A and Supplementary Table S2). Amino acid catabolism is regulated in response to nutrient availability, with TORC1 playing a central role in coordinating nitrogen and amino acid metabolism with cell growth (44, 45). This coordination is particularly relevant to fermentation, during which nutrient utilization must be balanced with rapid energy production. Consistent with this relationship, previous studies have demonstrated that TORC1 activity is required for efficient alcoholic fermentation and that reduced TORC1 activity markedly decreases fermentative capacity (46). More recently, deletion of the transcription factors Gcn4p, Gln3p, and Gat1p or the Npr1p kinase, all of which participate in the regulation of nitrogen and amino acid metabolism, was reported to enhance fermentation activity (47). These findings further support a close association between amino acid metabolism and fermentative performance. Curcumin has also been reported to inhibit TORC1 activity (48), which may appear to conflict with the established importance of TORC1 activity for efficient fermentation. However, in the present study, the intracellular levels of TCA-cycle intermediates and key amino acids associated with nitrogen metabolism tended to be lower in curcumin-treated cells than in untreated cells at 6 h after the initiation of fermentation (Fig. 7B). When considered together, these observations raise the possibility that, during the early phase of fermentation, curcumin transiently promotes amino acid utilization or catabolism in association with reduced TORC1 activity. In contrast, from the middle phase of fermentation onward, the expression of amino acid catabolism-related genes decreased and related intracellular metabolites accumulated, suggesting that amino acid catabolism was subsequently suppressed as the cells transitioned toward a metabolic state favorable for maintaining fermentation. Thus, the effects of curcumin on amino acid metabolism may not remain constant but may change as fermentation progresses.

The stress-adaptive and metabolic responses observed in curcumin-treated cells may not be entirely independent. Previous work has shown that, under osmotic stress conditions, Hog1 modulates TORC1 signaling and its downstream transcriptional programs by affecting the Sch9 and Tap42–PP2A branches (49). Thus, crosstalk between Hog1 and TORC1 may contribute to the coordination of stress adaptation and amino acid metabolism in curcumin-treated cells. However, whether the induction of stress adaptation-related genes and the downregulation of amino acid catabolism-related genes observed in this study are both mediated through this crosstalk remains unclear. Curcumin may enhance Hog1-associated stress-adaptive responses while also regulating amino acid catabolism through additional mechanisms. Future time-course analyses of Hog1 and TORC1 activities throughout fermentation will be required to determine how these signaling and metabolic responses are connected.

Overall, our findings provide new insights into the ecological and industrial significance of spice-derived compounds in microbial systems. Further elucidation of the molecular basis underlying the fermentation-enhancing effects of these compounds may contribute to a better understanding of plant–microbe interactions and the broader metabolic networks operating in ecosystems. From an applied perspective, the use of natural compounds to improve fermentation efficiency may represent a promising strategy for the food and bioenergy industries. Future studies should identify the direct molecular targets of curcumin and piperine, examine their effects on signaling pathways other than the HOG pathway, and determine whether their fermentation-enhancing effects are conserved in other microbial species.

## Materials and Methods

### Yeast Strains

Yeast strains used in this study included *Saccharomyces cerevisiae* strains X2180, K701, S288C, Σ1278b, X2180-1A, BMA64-1A, BY4741, and the *hog1Δ* mutant obtained from the National BioResource Project (NBRP), Japan. The fission yeast *Schizosaccharomyces pombe* wild-type strain L972 was also used.

### Test Compounds and Commercial Spice Products

The test compounds and commercial spice products used in this study are listed in Supplementary Table S3, together with their suppliers, catalog numbers, CAS registry numbers, and physical forms. Commercial food-grade curry powder and turmeric powder were purchased from S&B Foods Inc. (Tokyo, Japan). Unless otherwise indicated, compounds and spice products supplied as powders were added directly to the fermentation medium without prior dissolution. Liquid compounds were added directly to the medium. The concentrations used in individual experiments are described in the corresponding experimental sections.

### Primers

Primers used in this study for RT-qPCR analysis were designed based on the coding sequences of target genes in *Saccharomyces cerevisiae* and synthesized by Thermo

Fisher Scientific K.K. The following primers (5’ to 3’) were used:

ACT1 (*Fw_scACT1_307_334*: ATTATATGTTTAGAGGTTGCTGCTTTGG, *Rv_scACT1_591_566*: CAATTCGTTGTAGAAGGTATGATGCC); HXT1 (*HXT1_1259_F*: CATCTTCAAAGGGTCGTGGT, *HXT1_1435_R*: TCAAGAAACCCAGATGCAG);

PAU9 (*PAU9_5_F*: TGACTGGTATTGCCCAGAC, *PAU9_108_R*: ACCGTGCCTTGATAGAGCAC);

DAN1 (*DAN1_136_F*: GGCGCCCTCATTTACTGAATA, *DAN1_338_R*: GCAAGTGCGGAATAGAC);

GPP2 (*GPP2_283_F*: CGTGACTGTAAGGTGTGCAA, *GPP2_476_R*: TCGCCTCTTCAGATA);

GPD1 (*GPD1_386_F*: TTTTGCCGTATCTGTAGC, *GPD1_627_R*: AGAAATTCACCTTTTG);

TFS1 (*TFS1_255_F*: CGACCGTTTCACACTGGG, *TFS1_432_R*: CCCCTTGTATTGA);

HSP12 (*HSP12_156_F*: CTTCCCAGGGTCCACGACT, *HSP12_323_R*: TTGTGTGGTCTTCTCACC).

Gene-specific expression levels were calculated using the ΔΔCt method with *ACT1* as an internal control.

### Media and preculture conditions

Unless otherwise noted, fermentation assays were performed using YPD20 medium (1% yeast extract, 2% peptone, and 20% glucose). Yeast cells at the stationary phase (24–36 hours after pre-culture) were used. *S. cerevisiae* was pre-cultured in YPD medium (1% yeast extract, 2% peptone, and 2% glucose), and *S. pombe* in YES medium (0.5 % yeast extract and 3% glucose).

### Tube-based Fermentation

Fermentation medium (5 mL) was dispensed into test tubes and inoculated with yeast at an OD_600_ of 0.1. For curcumin-treated samples, curcumin was added to a final concentration of 1.0 mg/mL. Untreated samples were used as controls. Test tubes were sealed with aluminum caps, and their initial weights were recorded. Cultures were incubated statically at 30°C in a water bath. At 24-hour intervals, condensation on the inner wall was wiped off, and the tube weight was measured again. CO_2_ production [g] was estimated from the weight loss and used to evaluate fermentation capacity.

### Fermograph-based Fermentation

Fermentation was also monitored using the AF-1101-10W Fermograph II-W (Atto Corporation), a gas-volume measuring system (50). A volume of 50 mL of fermentation medium was dispensed into sterile bottles and inoculated with yeast at an OD_600_ of 0.1. Curcumin was added at final concentrations ranging from 0.01 to 10 mg/mL; untreated samples served as controls. After sample tubes were set into the Fermograph device, fermentation was carried out statically at 30°C, and CO₂ gas production [L] was measured over time.

### Growth Assay

Growth assays were conducted under the same conditions as the fermentation assays. Yeast cells in the stationary phase were used. A volume of 50 mL of fermentation medium was dispensed into sterile bottles and inoculated to an OD_600_ of 0.1. Curcumin was added to a final concentration of 0.1 mg/mL for treated samples, with untreated controls. Bottles were sealed with aluminum foil and incubated statically at 30°C in a water bath. OD_600_ was measured every 12 hours.

### Ethanol concentration

After fermentation, culture samples were centrifuged to remove yeast cells, and the ethanol concentration in the resulting supernatant was determined enzymatically using an Enzytec Liquid Ethanol kit (E8340; AZmax Co., Ltd., Tokyo, Japan) according to the manufacturer’s instructions. Absorbance was measured at 340 nm. Ethanol concentrations were calculated after accounting for the dilution factor and are expressed as % (vol/vol).

### Transcriptome Analysis

Stationary-phase yeast cells were inoculated into 50 mL of YPD20 medium at an initial OD₆₀₀ of 0.1 and fermented statically at 30°C with or without curcumin (0.1 mg/mL). After 12 h of fermentation, cell pellets equivalent to an OD₆₀₀ of 40 were collected, and total RNA was extracted using a FastGene RNA Purification Kit (Nippon Genetics Co., Ltd., Tokyo, Japan). Two independent biological samples were prepared for each condition. RNA quality assessment, library preparation, and sequencing were performed by Rhelixa Inc. (Tokyo, Japan). Poly(A)-selected, strand-specific libraries were prepared and sequenced on an Illumina NovaSeq X Plus platform to generate 150-bp paired-end reads. The resulting sequence data were analyzed in-house to compare gene expression profiles between curcumin-treated and untreated cells. Gene Ontology (GO) enrichment analysis was performed separately for upregulated and downregulated genes.

### RT-qPCR Analysis

Yeast cells were cultured as described above. After 12 hours of fermentation, cell pellets equivalent to OD_600_ = 40 were collected. RNA extraction was performed using the FastGene™ RNA Purification Kit, and cDNA synthesis was carried out using the PrimeScript™ RT reagent Kit (Takara Bio Inc.). For primer mix, 5 µL each of forward and reverse primers (10 µM) were mixed with 90 µL of sterile water. PCR reactions (10 µL total volume) consisted of 5 µL PowerTrack SYBR Green Master Mix (Thermo Fisher Scientific), 2 µL primer mix, 2 µL cDNA, and 1 µL sterile water. RT-qPCR was conducted using a LightCycler 96 (Roche), and gene expression levels were quantified using the ΔΔCt method with *ACT1* as the internal control.

### Zymolyase Sensitivity Assay

To evaluate cell wall integrity, a Zymolyase sensitivity assay was performed. Wild-type cells (with or without curcumin treatment) and the *Δhog1* mutant were cultured under fermentation conditions at 30°C for 24 hours. After cultivation, cells were harvested and resuspended in buffer to an optical density of OD_600_ = 1.0 in a final volume of 4 mL. Zymolyase 100T (Nacalai Tesque) was freshly prepared at a concentration of 100 µg/mL. To initiate digestion, 1 mL of the Zymolyase solution was added to the cell suspension (final concentration: 20 µg/mL in 5 mL total volume). As a negative control, 1 mL of buffer without Zymolyase was added to separate tubes. All samples were incubated at 30°C with gentle shaking. Cell lysis was monitored by measuring OD_600_ at 30-minute intervals. A decrease in OD_600_ over time was interpreted as increased sensitivity to Zymolyase-mediated cell wall digestion.

### Metabolomic analysis and data processing

Metabolite extraction and metabolomic analysis were performed using a GCMS- QP2020 NX system coupled with GC-2030 Nexis (Shimadzu, Kyoto, Japan) according to a modified version of a previously published protocol (51). Yeast pellet samples were extracted with 300 µL of extraction solvent prepared as a 10:1 mixture of mass spectrometry grade 100% methanol and a ^13^C-ribitol stock solution (0.2 mg/mL). The samples were subjected to ultrasonication for 10 min. Subsequently, 200 µL of chloroform was added, and the mixture was gently mixed for approximately 5 s. Then, 400 µL of MS-grade water was added, followed by vortex for approximately 15 s. After mixing, the samples were centrifuged at 10,000 *g* for 10 min at 4°C. Of the three resulting phases, 600 µL of the upper polar phase was collected and transferred to a new 1.5 mL tube. The polar phase was further aliquoted into two new 1.5 mL tubes, with 200 µL allocated for GC-MS analysis. The aliquoted samples were evaporated to dryness using a centrifugal evaporator and stored at 4°C until further analysis.

The 200 µL aliquots prepared for GC-MS analysis were further dried for 30 min using a centrifugal evaporator. Methoximation was performed by adding 40 µL of methoxyamination reagent, consisting of 4-(dimethylamino) pyridine (5 mg/mL) and methoxyamine hydrochloride (40 mg/mL) dissolved in pyridine, followed by incubation at 30°C with shaking at 200 rpm for 90 min. Subsequently, 70 µL of BSTFA containing 1% TMCS was added, and trimethylsilylation was carried out at 37°C with shaking at 200 rpm for 45 min. After derivatization, 80 µL of the reaction mixture was transferred to a GC-MS vial for metabolite analysis. In parallel, derivatized metabolite mixture standards for GC and GCMS-based metabolomics, as well as an alkane standard (C_9_–C_40_ Qualitative Retention Time Index Standard), were analyzed to aid in peak annotation.

GC-MS analysis was performed under the conditions summarized in Table 1. The GC oven temperature program was as follows: the temperature was set at 100°C at 0 min, held at 100°C until 4 min, increased to 320°C by 26 min, and held at 320°C until 37 min.

Metabolite identities were verified by comparison with spectral libraries of authentic standards in the Golm Metabolome Database (52). Detected peaks were evaluated using Xcalibur 4.1 software (Thermo Fisher Scientific, USA).

## Data Availability

The RNA sequence data generated in this study have been deposited in the DDBJ Sequence Read Archive (DRA) under BioProject accession number PRJDB35914.

## Acknowledgments

We thank Asako Higuchi for her technical assistance. We are also grateful to Dr. Ryo Nasuno and the NBRP Japan for the strains.

## Author Contributions

Y.N. and D.W. designed research; Y.N., S.K., Y.M., K.S., M.W. and Y.S. performed research; Y.N., S.K., T.T., and T.S. analyzed data; and Y.N. wrote the paper.

