## Supplementary Information for "Curcumin Enhances Alcoholic Fermentation in Yeast by Promoting Osmotic Stress Adaptation"

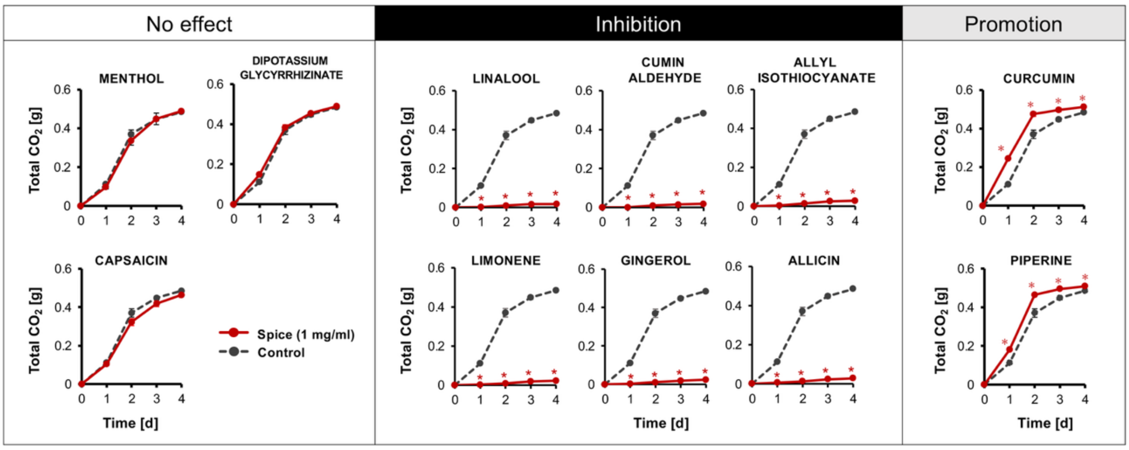


**Supplemental Figure S1. Effect of spice components on alcoholic fermentation.** There are three types of effects of the 11 spice components on alcoholic fermentation. Total $CO_2$ production (g) of strain X2180 in the presence or absence of 11 spice components (1 mg/mL). Data represent the mean ± standard deviation (SD) from three independent experiments. An asterisk indicates a significant difference from the untreated control at the corresponding time point (Student's *t* test, $P < 0.05$).

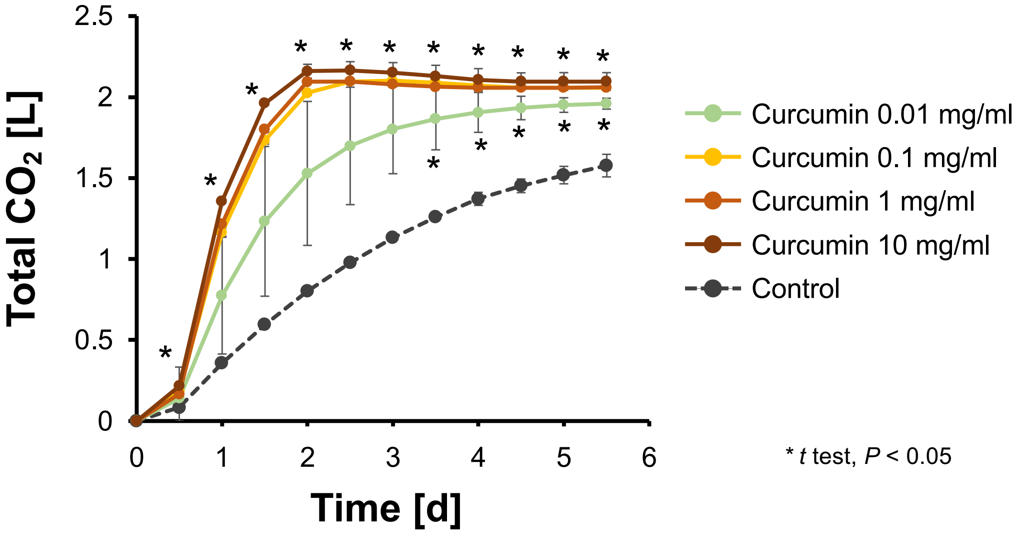


**Supplemental Figure S2. Effect of different concentrations of curcumin on alcoholic fermentation.**

Total $CO_2$ production (L) of strain X2180 in the presence or absence of curcumin (0.01, 0.1, 1, 10 mg/mL). Data represent the mean ± standard deviation (SD) from three independent experiments. An asterisk indicates a significant difference from the untreated control at the corresponding time point (Student's *t* test, $P < 0.05$).

| | | (A) Fold change at each time point (vs 0 h) | | | | | | | (B) Fold change (+ Curcumin vs Control at each time point) | | |
|---|---|---|---|---|---|---|---|---|---|---|---|
| | | Control | | | | + Curcumin | | | | | |
| | | 0h | 6h | 12h | 24h | 6h | 12h | 24h | 6h | 12h | 24h |
| Sugars | Glucose | 0.0 | 11.0* | 3.4* | 5.6* | 11.2* | 3.2* | 5.4* | 0.2 | -0.1 | -0.2 |
| | Fructose | 0.0 | 0.6 | 0.6* | 2.0* | -39* | 0.6 | 1.1 | -39.9 | 0.0 | -0.9 |
| | Sucrose | 0.0 | 1.7 | 0.0 | nd | 0.1 | -1.2 | nd | -1.6* | -1.2 | nd |
| | Maltose | 0.0 | 3.5 | 0.5 | -0.7 | 2.3* | -1.5 | nd | -1.1 | -2.0 | nd |
| | Galactose | nd | nd | nd | nd | nd | nd | nd | nd | -0.3 | -1.2 |
| | *β*-1,6-aGlc | nd | nd | nd | nd | nd | nd | nd | -0.8 | -0.1 | -0.4 |
| Sugar phosphate | G1P | 0.0 | nd | -2.6* | -0.6 | nd | -2.5* | -2.2* | nd | 0.0 | -1.6 |
| | G3P | 0.0 | 1.2* | 0.9 | 3.7* | -0.3 | 1.1 | 1.6* | -1.5* | 0.2 | -2.0 |
| Sugar alcohol | Ribitol | 0.0 | -0.3 | -0.4 | 0.0 | -1.6* | -0.5 | 0.4 | -1.3* | -0.1 | 0.4 |
| Cyclic sugar alcohol | Myo-Inositol | 0.0 | 2.4 | -0.8 | -0.7 | 2.2* | -1.2 | -1.3 | -0.2 | -0.4 | -0.6 |
| Saturated fatty acid | Palmitate | 0.0 | 0.5 | -1.4* | -1.0* | 0.0 | -1.0* | -1.0 | -0.5 | 0.3 | 0.1 |
| Glycolysis end product | Pyruvate | 0.0 | nd | nd | nd | nd | nd | nd | nd | nd | nd |
| TCA cycle | Citrate | 0.0 | -2.5* | -1.9* | -0.7 | -4.1* | -2.3* | 1.8* | -1.6* | -0.4 | 2.5* |
| | 2-Oxoglutarate | 0.0 | -0.9 | -0.9 | -2.6* | -1.6 | -0.9 | -0.5 | -0.7 | 0.0 | 2.1* |
| | Malate | 0.0 | -3.5* | -2.2* | -3.7* | -4.0* | -3.5* | -3.4* | -0.5 | -1.3* | 0.3 |
| Minor metabolite in yeast | Lactate | 0.0 | 2.6 | 2.1 | -0.9 | 6.4 | 5.1 | -2.2 | 3.8 | 3.0 | -1.3 |
| Aromatic alcohol | Benzylalcohol | nd | nd | nd | nd | nd | nd | nd | nd | nd | nd |
| Phospholipid component | Ethanolamine | 0.0 | 0.6 | 0.3 | 2.3* | nd | -0.7* | 1.6 | nd | -1.0 | -0.7 |
| Lipid metabolism related | Glycerol | 0.0 | 12.3* | 8.2* | 9.2* | 12.2* | 8.6* | 8.0* | 0.0 | 0.4 | -1.2* |
| Carbohydrate metabolism related | Glycerate | nd | nd | nd | nd | nd | nd | nd | -1.8 | nd | nd |
| Branched-Chain AAs | Valine | 0.0 | -2.4 | -1.2 | -4.4 | -3.1 | -2.4 | -3.6 | -0.7 | -1.2 | 0.8 |
| | Leucine | 0.0 | 1.0 | 0.3 | -1.4 | 1.5 | 0.3 | -1.8 | 0.5 | 0.1 | -0.4 |
| | Isoleucine | 0.0 | 1.9* | -2.8* | -3.4* | 1.7* | -1.5* | -4.1* | -0.2 | 1.3 | -0.6* |
| Central nitrogen metabolism and AA intermediates | Aspartate | 0.0 | 3.4* | 4.9* | 4.2* | 1.7* | 4.8* | 3.3* | -1.7 | 0.0 | -1.0 |
| | Asparagine | 0.0 | 0.3 | -0.6 | -0.8 | -1.4* | 0.1 | 0.9 | -1.7* | 0.6 | 1.7* |
| | Glutamate | 0.0 | -1.5* | -0.5 | -2.0* | -2.7* | -0.5 | -0.1 | -1.3* | 0.0 | 1.9* |
| | Glutamine | 0.0 | -0.3 | -1.9* | -2.7 | -1.4 | -1.8* | -1.4 | -1.0 | 0.0 | 1.3 |
| | Pyroglutamate | 0.0 | -1.5* | -0.5 | -2.1* | -2.8* | -0.5 | -0.1 | -1.3* | 0.0 | 1.9* |
| | Ornithine | 0.0 | -2.5* | -0.3 | -0.9 | -4.6* | -0.1 | -1.3* | -2.1* | 0.2 | -0.5 |
| | Arginine | 0.0 | 9.2* | -0.8 | -3.6* | 9.5* | -2.1* | nd | 0.3 | -1.2 | nd |
| | Lysine | 0.0 | 1.1 | 1.9* | 0.1 | 1.1* | 2.4* | -0.9* | 0.0 | 0.5 | -1.0 |
| | Homoserine | nd | nd | nd | nd | nd | nd | nd | nd | -0.6 | -0.8* |
| | *O*-Acetylserine | 0.0 | 40.0 | 39.3 | 38.9 | 39.9 | 38.2 | 39.4 | -0.1 | -1.1 | 0.4 |
| Polar AAs involved in sugar and one-carbon metabolism | Serine | 0.0 | 3.2* | 2.0 | 1.0 | 1.7* | 1.9 | -3.1 | -1.5* | 0.0 | -4.1* |
| | Threonine | 0.0 | 5.5* | 6.2* | 4.3* | 4.3* | 6.1* | 1.7 | -1.2* | -0.1 | -2.6 |
| | Methionine | 0.0 | nd | 1.4* | 0.8 | nd | 2.2* | 0.0 | nd | 0.8 | -0.9 |
| | Threonate | 0.0 | 3.0* | 4.6* | 3.9* | 1.2* | 4.6* | 2.8* | -1.9* | 0.0 | -1.1 |
| AAs related to cellular structure and stress response | Glycine | 0.0 | 2.8* | 2.8 | 2.3* | 2.9* | 2.5* | 3.4* | 0.1 | -0.3 | 1.1 |
| | Proline | 0.0 | 2.0* | 0.1 | -0.8* | 0.6 | 0.6 | -0.8 | -1.4 | 0.5 | 0.0 |
| Aromatic AAs | Phenylalanine | 0.0 | -1.6* | -1.0 | -2.6* | -2.8 | -1.0 | -0.6 | -1.2* | 0.0 | 2.0 |
| | Tyrosine | 0.0 | 1.5 | 0.6 | 0.1 | 1.0* | -0.5 | 0.3 | -0.6 | -1.1 | 0.2 |
| | Tryptophan | 0.0 | 0.5 | -0.3 | -1.2* | 0.3 | -1.6* | -1.4 | -0.3 | -1.2 | -0.2 |
| Trp derivative | Kynurenic acid | 0.0 | 0.1 | 0.4* | 0.1 | -0.1 | -0.1 | 0.4 | -0.1 | -0.5 | 0.3 |

Log2 fold change: 5, 0, -5

**Supplemental Figure S3. Heat map of intracellular metabolites during alcoholic fermentation with or without curcumin.**

Samples were collected at 0, 6, 12, and 24 h after the start of fermentation, and metabolite levels are shown as relative abundances normalized to cell biomass. (A) Log2 fold change at each time point versus 0 h shows the change of metabolite levels relative to 0 h, and (B) log2 fold change in curcumin-treated cells versus control at each time point shows the effect of curcumin. Student's *t* test was performed for each comparison ((A) *vs* 0 h; (B) *vs* control at each time point), and * indicates $P < 0.05$. Abbreviations: AA (amino acid), *β*-1,6-aGlc (*β*-1,6-anhydroglucose), G1P (glucose-1-phosphate), G3P (glycerol-3-phosphate).

**Supplemental Table S1. List of genes upregulated in the presence of curcumin compared with no curcumin.**

| Gene_ID | Gene name | Description | log2 fold change |
|---|---|---|---|
| **YHR094C** | HXT1 | Low-affinity glucose transporter | 7.42 |
| **YBL108C-A** | PAU9 | Seripauperin family protein | 6.56 |
| **YHL046C** | PAU13 | Seripauperin family protein | 5.59 |
| **YJR150C** | DAN1 | Cell wall mannoprotein expressed under anaerobic conditions | 5.48 |
| **YAL068C** | PAU8 | Seripauperin family protein | 5.17 |
| **YCR102W-A** | YCR102W-A | Putative uncharacterized protein | 5.17 |
| **YPL095C** | EEB1 | Acyl-coenzyme A:ethanol O-acyltransferase | 5.05 |
| **YBR072W** | HSP26 | Small heat shock protein | 4.87 |
| **YOR394W** | PAU21 | Seripauperin family protein | 4.32 |
| **YLR109W** | AHP1 | Alkyl hydroperoxide reductase | 4.22 |
| **YBL107W-A** | YBL107W-A | Putative uncharacterized protein | 4.21 |
| **YBR301W** | PAU24 | Seripauperin family protein | 4.19 |
| **YCR102C** | YCR102C | Putative protein of unknown function | 4.06 |
| **YOR273C** | TPO4 | Polyamine transporter of the major facilitator superfamily | 4.01 |
| **YGR131W** | FHN1 | Putative plasma membrane protein | 3.98 |
| **YBL108W** | YBL108W | Putative uncharacterized protein | 3.91 |
| **YOL150C** | YOL150C | Putative uncharacterized protein | 3.84 |
| **YCR104W** | PAU3 | Seripauperin family protein | 3.75 |
| **YLR303W** | MET17 | O-acetylhomoserine/O-acetylserine sulfhydrylase | 3.69 |
| **YOL151W** | GRE2 | NADPH-dependent methylglyoxal reductase involved in stress response | 3.54 |
| **YDR542W** | PAU10 | Seripauperin family protein | 3.48 |
| **YGR088W** | CTT1 | Cytosolic catalase T | 3.46 |
| **YDR345C** | HXT3 | Low-affinity glucose transporter | 3.40 |

| | | | |
|---|---|---|---|
| **YNL134C** | YNL134C | Putative protein of unknown function | 3.15 |
| **YBR300C** | YBR300C | Putative uncharacterized protein | 3.13 |
| **YBR244W** | GPX2 | Phospholipid hydroperoxide glutathione peroxidase | 3.06 |
| **YGL157W** | ARI1 | NADPH-dependent aldehyde reductase | 2.93 |
| **YOR009W** | TIR4 | Cell wall mannoprotein induced by anaerobic conditions | 2.91 |
| **YGL263W** | COS12 | Member of the DUP380 family of subtelomeric proteins | 2.90 |
| **YOR225W** | YOR225W | Putative uncharacterized protein | 2.85 |
| **YER069W** | ARG5 | Acetylglutamate synthase and N-acetylglutamate kinase | 2.85 |
| **YLR286C** | CTS1 | Endochitinase involved in cell separation | 2.83 |
| **YOL002C** | IZH2 | Membrane protein involved in zinc homeostasis | 2.82 |
| **YNR067C** | DSE4 | Cell wall glucanase involved in daughter cell separation | 2.79 |
| **tD(GUC)N** | tD(GUC)N | Aspartic acid tRNA | 2.73 |
| **YNL327W** | EGT2 | GPI-anchored cell wall protein involved in cell separation | 2.73 |
| **YHR143W** | DSE2 | Cell wall protein involved in daughter cell separation | 2.68 |
| **YOR226C** | ISU2 | Mitochondrial iron-sulfur cluster assembly protein | 2.66 |
| **YJR073C** | OPI3 | Phospholipid methyltransferase | 2.62 |
| **YGL028C** | SCW11 | Cell wall protein with glucanase activity involved in cell separation | 2.58 |
| **YLR461W** | PAU4 | Seripauperin family protein | 2.50 |
| **YOL165C** | AAD15 | Putative aryl-alcohol dehydrogenase | 2.38 |
| **YER124C** | DSE1 | Cell wall protein involved in daughter cell separation | 2.36 |
| **YPL250C** | ATG41 | Autophagy-related protein | 2.32 |
| **YNL014W** | HEF3 | Translational elongation factor EF-3 family protein | 2.28 |

| | | | |
|---|---|---|---|
| **YER091C** | MET6 | 5-methyltetrahydrofolate-homocysteine methyltransferase | 2.28 |
| **YKR075W-A** | YKR075W-A | Putative uncharacterized protein | 2.24 |
| **YGR209C** | TRX2 | Cytoplasmic thioredoxin | 2.23 |
| **YAL061W** | BDH2 | Putative (R,R)-butanediol dehydrogenase | 2.23 |
| **YER062C** | GPP2 | Glycerol-1-phosphatase involved in glycerol biosynthesis | 2.21 |
| **YPL256C** | CLN2 | G1/S-specific cyclin | 2.18 |
| **YKL001C** | MET14 | Adenylylsulfate kinase | 2.17 |
| **YKR075C** | YKR075C | Putative protein of unknown function | 2.15 |
| **YNL195C** | YNL195C | Putative protein of unknown function | 2.12 |
| **YBR071W** | YBR071W | Putative protein of unknown function | 2.11 |
| **YKL163W** | PIR3 | Cell wall protein of the PIR family | 2.06 |
| **YML131W** | YML131W | Putative protein of unknown function | 2.03 |
| **YOL101C** | IZH4 | Membrane protein involved in zinc homeostasis | 2.02 |

**Supplemental Table S2. List of genes downregulated in the presence of curcumin compared with no curcumin.**

| Gene_ID | Gene name | Description | $log_2$ fold change |
|---|---|---|---|
| **YLR307C-A** | DPA10 | Putative mitochondrial protein | -7.64 |
| **YHR126C** | ANS1 | Putative GPI protein | -6.23 |
| **YLR307W** | CDA1 | Chitin deacetylase | -6.13 |
| **YOR314W** | YOR314W | Putative protein of unknown function | -5.79 |
| **YKR034W** | DAL80 | GATA-family transcriptional repressor regulating nitrogen catabolic genes | -4.71 |
| **YKL177W** | YKL177W | Dubious ORF overlapping STE3 | -4.67 |
| **YKR033C** | YKR033C | Dubious ORF overlapping DAL80 | -4.56 |
| **YBR296C** | PHO89 | Plasma membrane Na+/Pi cotransporter | -4.43 |
| **YOR313C** | SPS4 | Protein whose expression is induced during sporulation | -4.26 |
| **YKL178C** | STE3 | Receptor for a factor pheromone | -4.08 |
| **YDR119W-A** | COX26 | Subunit of cytochrome C oxidase complex | -4.07 |
| **YHR137C-A** | YHR137C-A | Dubious ORF overlapping ARO9/YHR137W | -4.06 |
| **YHR137W** | ARO9 | Aromatic aminotransferase II | -4.00 |
| **YFR023W** | PES4 | Putative RNA binding protein | -3.98 |
| **YLR365W** | YLR365W | Putative conserved nonessential protein | -3.46 |
| **YJR078W** | BNA2 | Tryptophan 2,3-dioxygenase or indoleamine 2,3-dioxygenase | -3.42 |
| **snR64** | snR64 | C/D box small nucleolar RNA | -3.35 |
| **YDR380W** | ARO10 | Phenylpyruvate decarboxylase | -3.33 |
| **YHR015W** | MIP6 | mRNA-binding protein | -3.31 |
| **YLR438W** | CAR2 | L-ornithine transaminase | -3.30 |
| **YLR437C-A** | YLR437C-A | Dubious ORF overlapping CAR2/YLR438W | -3.28 |
| **YDR342C** | HXT7 | High-affinity glucose transporter | -3.23 |
| **YOR214C** | SPR2 | Putative spore wall protein | -3.20 |
| **YPL111W** | CAR1 | Arginase, catabolizes arginine to ornithine and urea | -3.07 |

| | | | |
|---|---|---|---|
| **YDR042C** | YDR042C | Uncharacterized protein upregulated in the ssu72-ts69 mutant | -2.88 |
| **snR128** | snR128 | C/D box small nucleolar RNA | -2.88 |
| **YHR214C-E** | YHR214C-E | Uncharacterized protein | -2.81 |
| **YHR092C** | HXT4 | High-affinity glucose transporter | -2.80 |
| **YBR200W-A** | YBR200W-A | Putative uncharacterized protein | -2.61 |
| **YBL098W** | BNA4 | Kynurenine 3-monooxygenase | -2.61 |
| **YLR296W** | YLR296W | Putative conserved uncharacterized protein | -2.50 |
| **YLR410W-B** | YLR410W-B | Retrotransposon TYA Gag and TYB Pol genes | -2.50 |
| **YOR315W** | SFG1 | Putative transcription factor regulating pseudohyphal and invasive growth | -2.46 |
| **YGR108W** | CLB1 | B-type cyclin | -2.44 |
| **YGR230W** | BNS1 | FEAR network component regulating Cdc14p release | -2.43 |
| **snR48** | snR48 | C/D box small nucleolar RNA | -2.38 |
| **YIL057C** | RGI2 | Uncharacterized protein involved in respiratory energy metabolism | -2.34 |
| **YOR186W** | YOR186W | Uncharacterized heat stress–responsive protein | -2.34 |
| **YKL102C** | YKL102C | Putative conserved oxidoreductase involved in citric acid tolerance | -2.32 |
| **YPR119W** | CLB2 | B-type cyclin | -2.32 |
| **YPR194C** | OPT2 | Oligopeptide transporter | -2.30 |
| **YBR297W** | MAL33 | MAL-activator protein | -2.28 |
| **YDL227C** | HO | Site-specific endonuclease; required for gene conversion at the MAT locus | -2.27 |
| **YHR184W** | SSP1 | BAR domain protein involved in the control of meiotic nuclear division | -2.26 |
| **YCL026C-A** | FRM2 | Type II NADH-dependent nitroreductase involved in stress responses | -2.25 |
| **YPL121C** | MEI5 | Meiosis-specific protein involved in meiotic recombination | -2.25 |
| **YDR523C** | SPS1 | STE20-family GCKIII protein kinase | -2.24 |

| | | | |
|---|---|---|---|
| **YJL037W** | IRC18 | Protein involved in outer spore wall assembly | -2.20 |
| **YBR040W** | FIG1 | Integral membrane protein required for efficient mating | -2.18 |
| **YDL181W** | INH1 | Protein that inhibits ATP hydrolysis by the F1F0-ATP synthase | -2.17 |
| **YPR123C** | YPR123C | Dubious ORF overlapping CTR | -2.16 |
| **YFL012W** | YFL012W | Uncharacterized sporulation-associated protein affecting rapamycin resistance | -2.15 |
| **YGR226C** | YGR226C | Dubious ORF overlapping AMA1/YGR225W | -2.13 |
| **YKL217W** | JEN1 | Monocarboxylate/proton symporter of the plasma membrane | -2.13 |
| **YGR177C** | ATF2 | Alcohol acetyltransferase | -2.11 |
| **YCR001W** | YCR001W | Putative conserved protein | -2.09 |
| **YIL162W** | SUC2 | Invertase | -2.08 |
| **YDR281C** | PHM6 | Phosphate-regulated uncharacterized protein | -2.08 |
| **YMR145C** | NDE1 | Mitochondrial external NADH dehydrogenase | -2.06 |
| **YPR124W** | CTR1 | High-affinity copper transporter of plasma membrane | -2.02 |

**Supplemental Table S3. Test compounds and commercial spice products used in this study.**

| Test compound or spice product | Supplier | Catalog no. | CAS RN | Form |
|---|---|---|---|---|
| ***Curcumin*** | Tokyo Chemical Industry Co., Ltd. | C0434 | 458-37-7 | Powder |
| ***Piperine*** | Tokyo Chemical Industry Co., Ltd. | P0460 | 94-62-2 | Powder |
| ***Menthol*** | Nacalai Tesque, Inc. | 21423-75 | 2216-51-5 | Powder |
| ***Capsaicin*** | Nacalai Tesque, Inc. | 07127-21 | 404-86-4 | Powder |
| ***Dipotassium glycyrrhizinate*** | FUJIFILM Wako Pure Chemical Corporation | 072-03862 | 68797-35-3 | Powder |
| ***Linalool*** | Nacalai Tesque, Inc. | 20505-22 | 78-70-6 | Liquid |
| ***Cuminaldehyde*** | Tokyo Chemical Industry Co., Ltd. | I0168 | 122-03-2 | Liquid |
| ***Limonene*** | Tokyo Chemical Industry Co., Ltd. | L0046 | 138-86-3 | Liquid |
| ***Gingerol*** | FUJIFILM Wako Pure Chemical Corporation | AG0035YB | 23513-14-6 | Powder |
| ***Allyl isothiocyanate*** | FUJIFILM Wako Pure Chemical Corporation | 0001317 | 57-06-7 | Liquid |
| ***Allicin*** | MedChemExpress | HY-N0315 | 539-86-6 | Liquid |
| ***Ferulic acid*** | Tokyo Chemical Industry Co., Ltd. | H0267 | 537-98-4 | Powder |
| ***Vanillin*** | FUJIFILM Wako Pure Chemical Corporation | 224-00682 | 121-33-5 | Powder |
| ***Demethoxycurcumin*** | Tokyo Chemical Industry Co., Ltd. | D6479 | 22608-11-3 | Powder |
| ***Bisdemethoxycurcumin*** | Tokyo Chemical Industry Co., Ltd. | B6548 | 33171-05-0 | Powder |
| ***Tetrahydrocurcumin*** | Tokyo Chemical Industry Co., Ltd. | T3587 | 36062-04-1 | Powder |

| | | | | |
|---|---|---|---|---|
| ***curry powder*** | S&B Foods Inc. | N/A | N/A | Powder |
| ***turmeric powder*** | S&B Foods Inc. | N/A | N/A | Powder |
